# Isolation of small extracellular vesicles from regenerating muscle tissue using Tangential Flow Filtration and Size Exclusion Chromatography

**DOI:** 10.1101/2024.02.14.580358

**Authors:** Uxia Gurriaran-Rodriguez, Michael A. Rudnicki

## Abstract

We have recently made the strikingly discovery that upon a muscle injury, Wnt7a is upregulated and secreted from new regenerating myofibers on the surface of exosomes to elicit its myogenerative response distally. Despite recent advances in extracellular vesicle (EVs) isolation from diverse tissues, there is still a lack of specific methodology to purify EVs from muscle tissue. To eliminate contamination with non-EV secreted proteins and cytoplasmic fragments, which are typically found when using classical methodology, such as ultracentrifugation, we adapted a protocol combining Tangential Flow Filtration (TFF) and Size Exclusion Chromatography (SEC). We found that this approach allows simultaneous purification of Wnt7a, bound to EVs (retentate fraction) and free non-EV Wnt7a (permeate fraction). Here we described this optimized protocol designed to specifically isolate EVs from hind limb muscle explants, without cross-contamination with other sources of non-EV bounded proteins. The first step of the protocol is to remove large EVs with sequential centrifugation. Extracellular vesicles are then concentrated and washed in exchange buffer by TFF. Lastly, SEC is performed to remove any soluble protein traces remaining after TFF. Overall, this procedure can be used to isolate EVs from conditioned media or biofluid that contains EVs derived from any cell type or tissue, improving reproducibility, efficiency, and purity of EVs preparations. Our purification protocol results in high purity EVs that maintain structural integrity and thus fully compatible with *in vitro* and *in vivo* bioactivity and analytic assays.

## INTRODUCTION

In the past decade, protein secretion mediated by extracellular vesicles (EVs), has emerged as a physiopathological relevant process for distal intercellular communication [1]. Exosomes represent a population of small EVs of endocytic origin involved in intercellular interaction through the transport of essential cargoes such DNA, RNA, or proteins to distal places [2]. Numerous studies have shown the potential of EVs-based therapies as nanocarriers to treat different pathologies [3,4]. Accordingly different techniques have emerged as standardized protocols to isolate EVs from different cell-derived conditioned media or biofluids [5–7] in agreement with the Minimal Information for Studies of Extracellular Vesicles (MISEV) [8,9]. Recently, tissue-derived EVs have been recognized as powerful role players in physiopathological process compared to biofluid-derived EVs, given the information enrichment from the tissue microenvironment [10,11]. Despite the extensive research in the field, the clinical implementation of these findings is still far from being translated, mainly due to the technical complexity of EVs isolation and administration for reproducible and reliable experimentation [12,13].

Particularly, there is an overwhelming evidence for the therapeutic use of EVs in muscle related pathologies [14–21]. In line with this, Wnt proteins, a conserved family of secreted glycoproteins that govern essential developmental, growth and regenerative stem cell processes, are a representative example of long-distance signaling and myoregenerative potential [22]. We have shown how Wnt7a/Frizzled signaling axis plays an essential role on myogenesis and muscle stem cell homeostasis at different levels [23–27]. Deciphering how Wnt7a release mechanisms are regulated in muscle is key for the therapeutic translation of all those discoveries and the implementation of standardized protocols for muscle research. Intriguingly, despite the relative hydrophobicity due to palmitoylation[28], in addition to acting in endocrine and paracrine signaling, Wnt proteins also have been suggested to activate signaling distally through several potential mechanisms such as lipoproteins, cytonemes, and association with soluble Wnt-binding proteins [29]. Recently, several studies have found that Wnt proteins act distally after secretion on EVs [30–32]. Indeed, we have recently discovered that upon an injury Wnt7a is secreted on the surface of EVs. More specifically, we have elucidated the Wnt7a structural motif that is required to direct Wnt7a onto EVs for long-range signaling, and its relevance in the myoregenerative response [33].

We identified the molecular sequence, within the Wnt7a structure, required for EV secretion, which we term the Exosome Binding Peptide (EBP). Mutation of EBP causes a disruption on long-range Wnt7a-EV secretion and a shift towards Non-EV bound Wnt7a secretion. This unprecedented mechanism is mediated by interaction with Coatomer proteins and independent of palmitoylation or WLS [34], the transmembrane protein that regulates Wnt classical secretion [33,35]. Notably, a single intravenously injection of myotube-derived EVs bearing Wnt7a [35] rescued muscle regeneration in Wnt7a-deficient dystrophic *mdx* mice.

To date, previous isolation protocols for muscle-derived EVs were designed for EV isolation from conditioned media from myogenic cells [36,37], or EVs derived from tissue after enzymatic digestion [38]. Here, we describe a detailed protocol to isolate small EVs secreted from regenerating muscle tissue explants combining Tangential Flow Filtration (TFF) and Size Exclusion Chromatography (SEC). TFF is an ultrafiltration technique that allows the separation of EVs based on size [39,40]. SEC provides a final purification step to separate free-protein cross-contaminants [41]. Combination these two techniques allow purification of EVs free of soluble proteins while maintaining structural integrity of EVs for downstream applications with an overall improved yield efficiency.

## RESULTS

### Tangential Flow Filtration purifies high-efficient EVs without soluble protein cross-contamination in cell-derived conditioned media

Conditioned media from Wnt7a-transfected HEK293T cells has shown that Wnt7a proteins are secreted on EVs surface, but also as non-EV bound free protein [33]. Here we compared the suitability of the golden-standard procedure for EVs purification, ultracentrifugation[42], with TFF to isolate Wnt7a-EVs from HEK293T cells. Ultracentrifugation purification of Wnt7a-transfected HEK293T conditioned media results in non-EV bound Wnt7a pellet down with EVs, producing a cross contamination difficult to discriminate (Fig. 1A). Conversely, TFF allows the physical separation of Wnt7a non-EV bound free protein that filtrates through the pores of the 500KDa cartridge and is collected in the permeate fraction, while Wnt7a-EVs are retained inside the cartridge as the retentate fraction (Fig. 1B). Noteworthy, the double lipid membrane was not affected upon isolation using TFF (Fig. 1C), compared to ultracentrifugation that can easily disrupt and burst the fragile EVs (Fig. 1D). Overall, the use of TFF results in increased yields with cleaner intact EVs compared to ultracentrifugation (Fig. 1E-F).

**Fig. 1.**
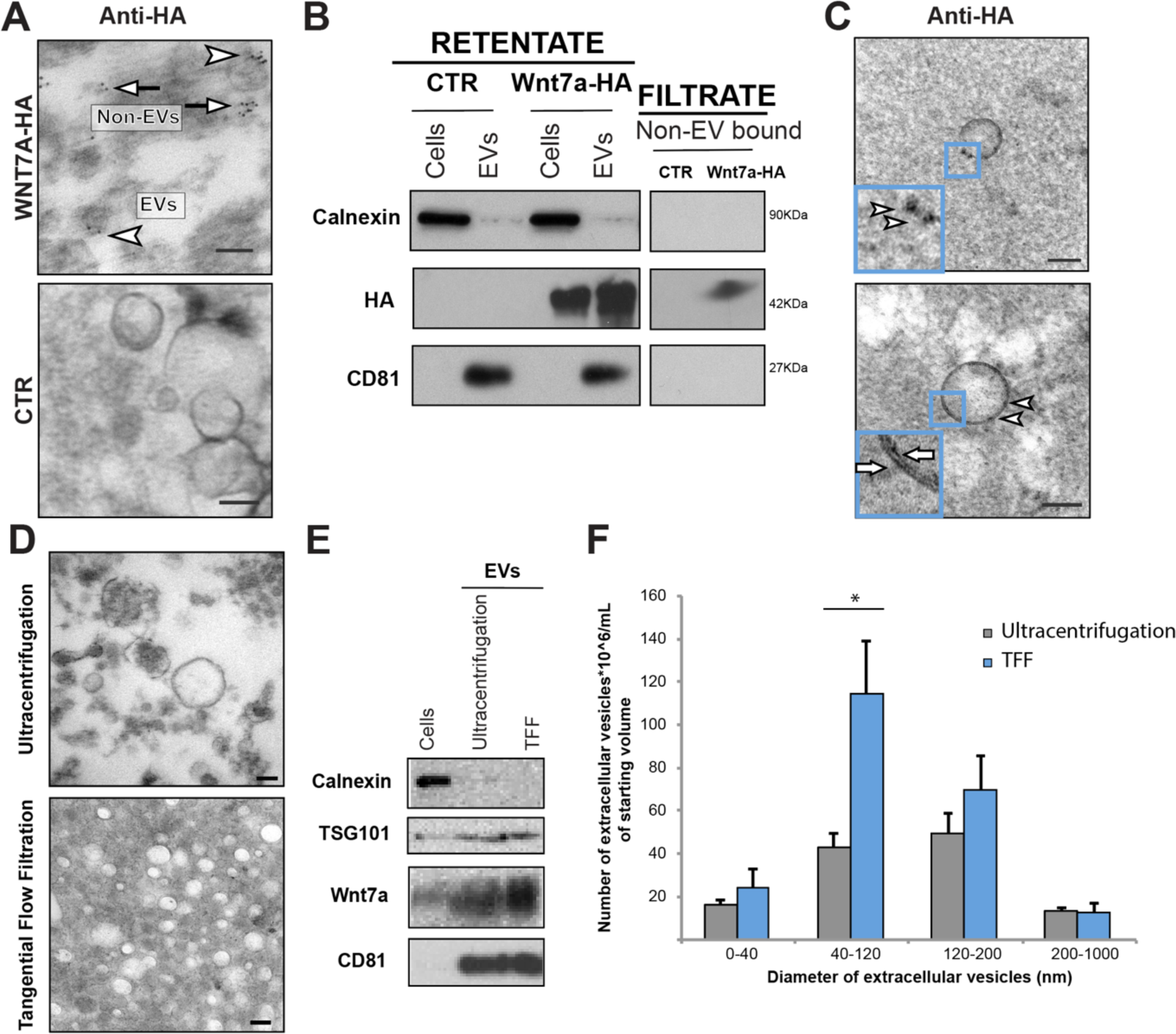
Tangential Flow Filtration allows purification of small EVs free of non-EV bound components. **A** iTEM of anti-HA immunogold labeling after ultracentrifugation protocol from Wnt7a-HA transfected HEK293T cells conditioned media showing isolation of non-EV bound Wnt7a cross-contamination (arrows) with Wnt7a-EVs (arrow heads)**. B** Immunoblot analysis for Wnt7a showing separation of non-EV bound Wnt7a (filtrate) vs Wnt7a-EVs (retentate) after TFF protocol. **C** iTEM of anti-HA immunogold labeling after TFF protocol from Wnt7a-HA transfected HEK293T cells conditioned media showing the absence of Wnt7a non-EV bound cross-contamination. Membrane bilayer indicated with arrows. **D** Comparative TEM of EVs isolated with ultracentrifugation and TFF**. E** Immunoblot analysis comparing EVs isolation efficiency with ultracentrifugation and TFF. **F** Comparative size distribution analysis of EVs samples after TFF protocol (blue bars) versus classical ultracentrifugation (grey bars). Scale bar 100nm. Empty plasmid (CTR), (n=3, mean ± s.e.m., *p<0.05).

### Muscle injury triggers release of muscle-derived extracellular vesicles

Transmission Electron Microscopy (TEM) revealed that 96h following injury, induced by cardiotoxin injection, new muscle regenerating myofibers release high numbers of multivesicular bodies (MBVs) that are filled with EVs destined for secretion. Likewise, the surrounding intersttium in the injured muscle is fully infiltrated with the secreted EVs. Conversely, in uninjured healthy muscle, infiltration of EVs cannot be detected (Fig. 2A). Inmmunogold labeling of new regenerating myofibers showed how Wnt7a is secreted on EVs surface from new regenerating myofibers that produce MVBs containing those Wnt7a-expressing EVs (Fig. 2B). Importantly, the same type of Wnt7a-expressing EVs were found in the surrounding injured tissue (Fig. 2C). This data suggests that new regenerating myofibers released Wnt7a on EVs to trigger a distal regenerative response. Here we present a specific protocol to isolate those released muscle-tissue derived EVs combining centrifugation, Tangential Flow Filtration (TFF) and Size Exclusion Chromatography (SEC) (Fig. 2D).

**Fig. 2.**
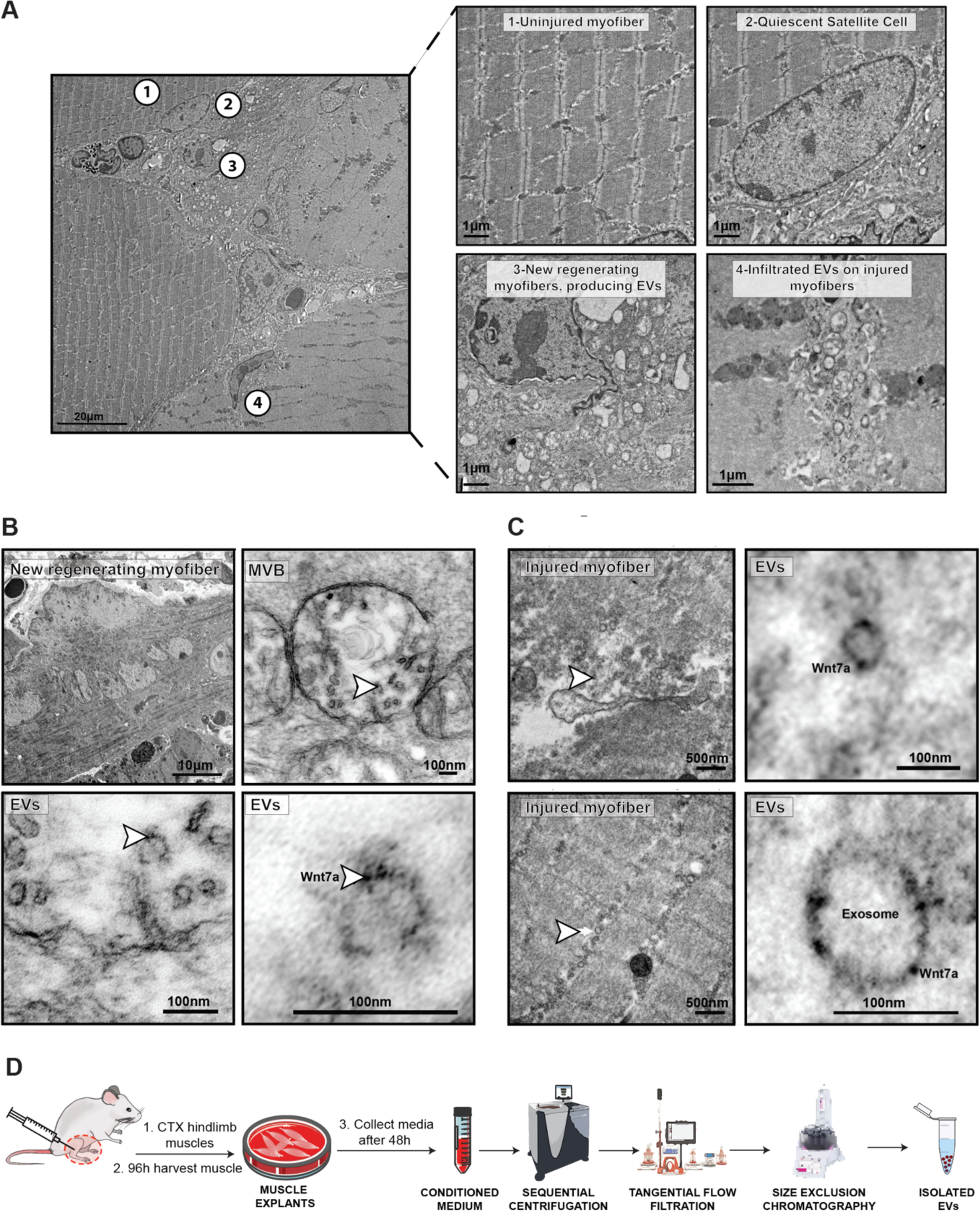
New regenerating myofibers secrete exosomes that target injured muscle. **A** Representative TEM image of a muscle transversal projection showing exosomes secretion 96 hours post injury. Uninjured muscle (1) containing quiescent satellite cells (2), does not secrete or uptake exosomes. Conversely, new regenerating myofibers secrete exosomes (3) infiltrating distal injured muscle to exert a myoregenerative response (4). **B** Representative anti-Wnt7a iTEM labeling of a new regenerative myofiber transversal projection showing MVBs containing Wnt7a-expressing exosomes 96 hours post injury. **C** Representative anti-Wnt7a iTEM labeling of an injured myofiber showing Wnt7a-expressing exosomes infiltration 96 hours post injury. Transmission Electron Microscopy (TEM), Multi Vesicular Body (MVB). **D** Workflow protocol of small EVs isolation derived from muscle injured tissue explants.

## EV PURIFICATION PROTOCOL

No pre-acquired expertise is needed more than being qualified to work with animals to obtained the tissue explants. All the other expertise required for this protocol could be easily achieved by a single person familiarized with bench work in a research lab by following this protocol. The entire procedure should be performed under sterile conditions, autoclaving all the non-disposable material but cartridges and pressure sensors could be damaged by the high temperatures. Given that this last material cannot be autoclaved is highly recommendable to install the TFF machine inside of a tissue culture hood to minimize contamination due to the high content of biomaterial and the porous nature of the cartridges where impurities can lead to contamination easily. Also, the required rotors for pre-clarification and last pellet down steps can be difficult to find when there are not several different rotors available. However, in the present protocol, several options have been exposed to make it easier for the researcher.

### Mice

This protocol has been standardized for muscle derived-EVs isolation with mice from different backgrounds, C57BL6 and C57BL/10ScSN. In our particular case, we have not found differences in terms of yield with different strains for these backgrounds. However, we cannot anticipate that changes in mouse strain could lead to differences in isolation yield. We recommend using mice aged 8-12 weeks independently of their sex, given that very young or old mice could have differences in EVs secretion. Housing, husbandry, and all experimental protocols for mice used in this study were performed in accordance with the guidelines established by the University of Ottawa Animal Care Committee, which is based on the guidelines of the Canadian Council on Animal Care.

### Reagents

All reagents were prepared and stored according to the manufacturer’s instructions.

The use of alternative reagents from other companies may not guarantee the anticipated results.

- Chlorohexidine, 2%
- Eye lubricator
- PBS without calcium and magnesium, pH 7.4 (Wisent, cat. no. 811-012-FL)
- FBS (Wisent, cat. no.098-150)
- Penicilyn/Streptomycin (Wisent, cat. no. 450-201-EL)
- DMEM media (Wisent, cat. no. 319-016-EL)
- Cardiotoxin, citotoxin I (Latoxan, cat. no. L81 02).
- NaOH (Fisher Scientific, cat. No. S318-500)
- Sodium Chloride, 0.9% (Baxter, cat. no. JB1301)
- MilliQ water
- Ice

### Equipment

- Anesthesia equipment
- Fur trimmer
- Insulin syringe, sterile 28 gauge (Fisher, cat. no. 36-104-1352)
- 3mL syringe (Plastipak; BD, cat. no. 309657)
- 22G × 1 1/2 needles, 0.7 mm × 40 mm (BD, cat. no. 305156)
- Fine scissors (Fine Science Tools, cat. no. 14060-11)
- Dissecting scissors (Fisherbrand, cat. no. 08-951-20)
- Spring scissors (Kent scientific, cat. no. INS600127)
- Forceps (Fisherbrand, cat. no. 10-270)
- Micro forceps
- Aluminum foil
- Styrofoam rack
- Water bath (Fisherbrand, model FSGPD05)
- 150mm sterile petri dishes
- Cell incubator
- TC hood
- Pipettes 50mL (Sarstedt, cat. no. 86.1685.001) and 25mL (Sarstedt, cat. no. 86.1689.001)
- 50-ml conical centrifuge tubes (Falcon, cat. no. 1495949A)
- 15-ml conical centrifuge tubes (Fisher, cat. no. 12565269)
- Thermo Scientific™ Sorvall™ Legend™ XT/XF Centrifuge with rotor (Fisher, cat. no. 75-333-839)
- Ultracentrifuge Optima L-90K (Beckman Coulter, cat. no. SKU 8043-30-1191)
- Rotor 45Ti (Beckman Coulter, cat. no. 339160)
- Polycarbonate bottle assembly, 70mL 38x 102 mm (Beckman Coulter, cat. no. 355622)
- Rotor JA-25.50 (Beckman Coulter, cat. no. 363058)
- Polycarbonate bottle with screw-on cap, 50mL 29x 104 mm (Beckman Coulter, cat. no. 357002)
- KrosFlo Research *2i* TFF System (Repligen, cat. no. SYR2-U20, ACSC-20K-0908 AND ACJR-U10)
- Pressure sensors (Repligen, cat. no. ACPM-799-01N)
- Hollow Fiber Cartdrige 500MWCO (Cytiva, cat. no UFP-500-E-H42LA)
- Tubing 14”diameter (SpectrumLabs, cat. no. ACTU-E14-25N).
- 1L Pyrex bottle
- Corning™ GL45 Three-Hole Delivery Cap (Fisher, cat. no. 10-462-474)
- Flexyboy with EVA Tubes & Luer Lock-50mL (Satorius Stedim North America, INC, cat. no FFB207567)
- pH paper strips (Fisher, cat. no. M10953330001)
- Becnhtop ultracentrifuge Optima Max 130 000
- Rotor TLA-100.3 (Beckman Coulter, cat. no. 349490)
- Open-top thickwall polycarbonate tube, 3,2mL 13×56mm tubes for TLA100.3 rotor (Beckman Coulter, cat. no. 362305).
- Delrin Tube Adaptors 11mm diameter for TLA.110 (Beckman Coulter, cat. no. 355919)
- Polypropylene tube with snap-on cap, 1.5mL, 11×39mm (Beckman Coulter, cat. no. 357448)
- NTA equipment
- Stericup, 0.22-uM polyethersulfone (Millipore, cat. no. S2GPU11RE)
- Metal stand

## REAGENT SETUP

### Cardiotoxin

Resuspend under sterile conditions the cardiotoxin powder to have a stock solution 100 μM. Dilute 1 mg in 1.47 ml of in 0,9% Sodium Chloride. Aliquot and freeze samples at −20°C. Working solution is 10 μM in 0,9% Sodium Chloride. Working solution needs to be prepared freshly on a TC hood and unused solution should be discarded appropriately. Cardiotoxin is amyotoxin and should be handled with care. Protect the eyes and skin by wearing gloves and working in a biosafety hood.

### FBS EVs-depleted

Centrifuge FBS under sterile conditions at 100,000 G for 90min using Beckman Coulter Optima L-100XP Ultracentrifuge, Rotor 45Ti (Beckman Coulter, cat. no. 339160) and tubes (Beckman Coulter, cat. no. 355622). Upon centrifugation carefully transfer the EVs-depleted FBS and make 50 ml aliquots. Storage them until use at −20°C.

### PBS microparticles-depleted

Centrifuge PBS 1x at 100,000 G for 90min using Beckman Coulter Optima L-100XP Ultracentrifuge, Rotor 45Ti (Beckman Coulter, cat. no. 339160) and tubes (Beckman Coulter, cat. no. 355622). Upon centrifugation filter the PBS using a Stericup under sterile conditions and storage it 1L sterile glass bottles.

### NaOH

Prepare NaOH 0.5 M in MilliQ water. Filter the NaOH under under sterile conditions using a Stericup and storage it 1 L sterile glass bottles.

## EQUIPMENT SETUP

### Tubing circuit setup

1. Introduce a piece of tubing in the cap of each wash bottle and add plugs to the other three holes of the cup (Fig. 3A). We highly recommend doing separate tubing for each washing bottle to prevent cross contamination. Transparent tubing is used for regular setups and white tubing reserved it only for tubing used with the pump.
2. Plug white tubing using female and male ports connecting the cap of the washing bottle with the auxiliary pump (Fig. 3A-B).
3. Connect the end of the washing tubing port that comes from the auxiliary pump to a T-valve (Fig. 3B).
4. Remove red caps from the cartridge and plug the three pressure sensors (Fig. 3C).
5. Screw female cap into the bottom auxiliary port of the cartridge (Fig. 3C).
6. Plug a piece of white tubing, using female and male ports connected to the feed and retentate pressure sensors respectively (Fig. 3D-E).
7. Connect the T-valve to the white tubing coming from the retentate fraction (Fig. 3D-E).
8. Connect the last third port of the T-valve to the Flexboy bag and its clamps (Fig. 3D-E).
9. Connect the second port of the Flexboy bag to the white tubbing coming from the feed flow (Fig. 3D-E).
10. Connect a transparent tubing to the permeate pressure sensor and a Flexboy bag if we want to collect the permeate fraction containing the fraction that is smaller than 500KDa, for further applications. Otherwise, connect a waste container (Fig. 3D-E).

### New cartridge rinsing and sanitation procedure

Cartridges need to be rinsed before use as they are stored in glycerol. It is recommendable to perform the cartridge activation the day before the purification due to long duration of the procedure.

1 Warm up sterile MilliQ water up to 50°C using one of the below approaches. Control temperature closely because if the temperature is above 50°C it can damage the cartridge.

A) A heating magnetic stirrer placed beside the auxiliary pump.
B) A microwave or water bath at the beginning of the procedure. However, using this procedure water temperature will not be maintained along the entire cleaning process.
2 Connect the circuit set up to the TFF machine (see Fig. 3). Ensure clamps on the Flexboy bag are open and everything is properly connected to avoid damaging filter or sample loss due to high pressure (Fig. 3D-E).
3 Connect the sterile warmed MilliQ water bottle to the auxiliary pump washing port (Fig. 3D-E).
4 Switch on the TFF machine, the auxiliary pump, and the monitor. All pressure (feed, permeate and retentate) should be 0.0 psi in the display of the main pump.
5 Login in into the software. Spectrum Labs-→KR2i Data Collection. Set up the machine on the display to work counterclockwise and the main pump on M mode (manual).
6 Introduce all the cartridge information in the display before starting the procedure. Consider that by default the TFF software is pre-setup with Repligeńs cartridges details and we are using a cartridge from Cytiva. Thus, we have to introduce Cytivás technical information manually, so the machine can register properly the cartridge in use and record all the technical details during the procedure accordingly. Fiber count: 2, Fiber ID: 1, Fiber length: 110, Area: 73cm^2^ (see Fig. 3F).
7 Disconnect retentate flow from the continued circuit and connect retentate and permeate fluxes in the waste. Screw a plastic cap on the T-valve to cover the open end of the retentate port.
8 Set up the flow rate of the auxiliary pump at 20mL/min, to flush the warmed MilliQ water bottle, and the feed flow of the main pump at 20mL/min with the back pressure valve and T-valve open (see Fig. 3G).
9 Start the auxiliary pump and the main pump flows. Hold these settings for 5 minutes aprox. until pressure is stable and there is no bubble in the tubing.
10 Stop the flow. Remove the plastic cap on the T-valve and connect retentate flow for recirculation, back to the continued circuit.
11 Set up flow rate of the auxiliary pump at 3mL/min and feed flow speed to 10ml/min and initiate the flow. Pressure should be P_f_ ∼2-4 psi, P_r_ ∼1.2-1.5 psi and P_p_ 0 psi. Keep these settings until 500mL are permeated (see Fig. 3F-G).

**Fig. 3.**
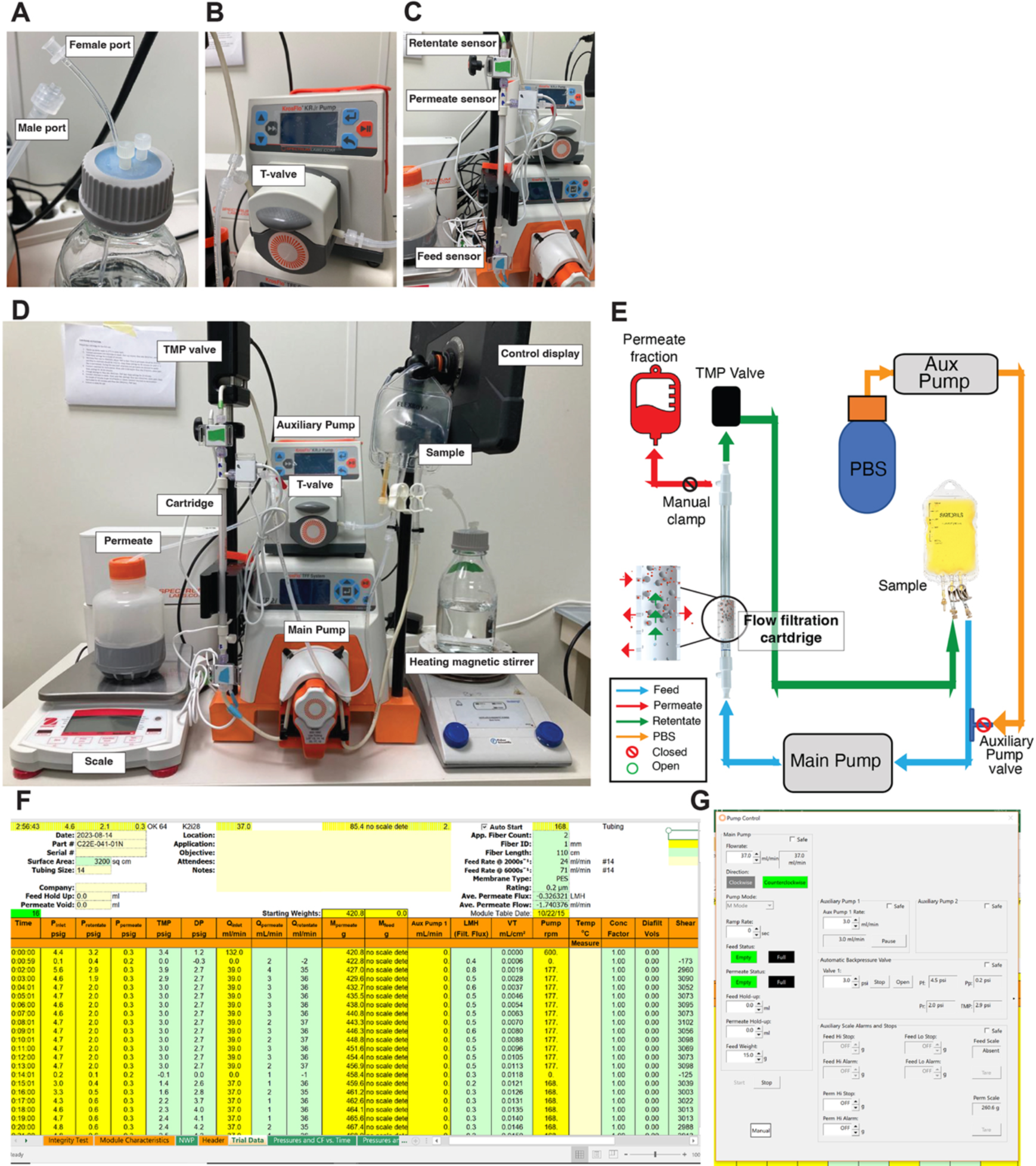
Tangential Flow Filtration setup. **A** Buffer exchange bottle showing female/male ports of the tubing circuit. **B** Auxiliary pump connected to the exchange/washing buffer showing special white tubing for the pump. **C** Hollow Fiber Cartridge with 500 MWCO connected to the Retentate/Permeate and Feed pressure sensors. **D** Entire TFF set up containing the machine and the circulation circuit with all the necessary components**. E** Representative schematic set-up of the TFF circuit. **F** Representative picture of the recording display measuring the different parameters along the purification TFF process. **G** Representative picture of the pre-settings pump control for the purification TFF process.

**Critical step.** Set up a permeate scale alarm in the settings panel to automatically stop the flow when a permeate volume of 500 ml is reached. Make sure to tare the scale before starting the procedure (Fig. 3D).

**Critical step.** Verify there is no leaking in the tubing or port connectors and adjust TMP to 5psi in the display. Once 5psi TMP is reached flow to permeate should be 9ml/min and flow to retentate should be 1ml/min. Verify this by checking Q inlet (ml/min), Q permeate (mL/min) and Q retentate (mL/min).

**Critical step.** If volume in the Flexboy bag is not steady, please modify flow rate of the auxiliary pump. If it is too low, bubbles will appear in the circuit, then increase flow rate, up to 3mL/min maximum. If it is too high, decrease up to 0.5mL/min. Thus, the volume entering the Flexboy bag is equal to the volume that is flowing through the cartridge.

12 Stop the machine and interchange the water bottle with a bottle containing sterile NaOH 0.5M to sanitize the circuit.
13 Set up the auxiliary pump at a flow rate of 1ml/min and the main pump at 37ml/min with the back pressure valve open to facilitate longer recirculation of NaOH.
14 Flux up to 100 ml of NaOH to sanitize the entire circuit. Set up a permeate scale alarm in the settings panel to automatically stop the flow when you have a permeate volume of 100 ml is reached.

**Critical step.** Keep the volume in the Flexboy bag steady if not proceed as in step 11.

15 Stop the flow and connect retentate and permeate to waste. Screw a plastic cap on the T-valve to cover the open end of the retentate port. Connect the sterile PBS bottle to the auxiliary pump instead of the NaOH bottle. Open auxiliary pump and T-valve so circuit is open to PBS and tubing is not obstructed. Make sure that back pressure valve is opened and and main pump is set up at 132 ml/min (see Fig. 3D-E).

**Critical step.** Given the high flux of the main pump (132 ml/min), there is no need to use the auxiliary pump to inject the PBS into the feed flow. Leaving all the ports open, the inherent high flow of the main pump will be sufficient for the washing PBS to get to the circuit.

16 Wash with 50 ml of PBS at maximum speed, 132 ml/min, to flux out remaining NaOH. Stop the flow. Remove the plastic cap on the T-valve and connect retentate flow for recirculation. Initiate the flux with a speed rate of 37ml/min and TMP 3psi.

**Critical step.** Recirculate 50 ml of PBS and check with pH paper strips if permeate fraction pH decreased up to 7, meaning non traces of NaOH remain inside the circuit that could damage the upcoming sample purification. Otherwise keep recirculating PBS until optimal neutral pH 7 has been reached.

17 Column is ready for use.

**Pause point.** If sample filtration is going to be performed the following day, we can leave all the circuit set up already installed in the machine. Otherwise, remove it from the machine and store it in a safe place. The entire circuit including cartridge, tubing and pressure sensors can be stored all interconnected (see Fig. 3D-E).

**Critical step.** Never store the cartridge and sensors in NaOH 0.5 M, as the material can be damaged as well as difficult to disconnected upon prolonged exposures to NaOH.

## PROCEDURE

### Sample preparation

1 Bring mice into the surgical room and inject analgesia accordingly to the animal procedure protocol of the institution (p.e. subcutaneous 0.1 mg/kg buprenorphine 4h prior to surgical procedure).

**Critical step.** Is not recommended to perform more than 4 mice per experiment as it would be difficult to handle.

2 Anesthetize the mice with isoflurane (p.e. 1L/min, 2% isofluorane, or equivalent) for cardiotoxin intramuscular injection. Apply eye lubrication gel in the ocular area to avoid eye damage due to dryness, as anesthetized mice leave their eyes open. Inject 1 ml of saline subcutaneous to avoid dehydration. Shave both lower hind limbs and wipe the area with chlorohexidine to clean the area.

**Critical step.** Do not proceed to step 3 until you verify there are no signs of reactivity upon gentle pressure on the toes with a surgical forceps.

3 Place the mouse in supine position and inject CTX solution intramuscularly using a sterile 28-gauge insulin syringe. Insert the needle with the bevel downward through the ankle parallel to the tibia until it reaches the knee (half of the needle length). Then inject the solution slowly and at the same time gradually remove the needle. On each leg inject 50 μl of Cardiotoxin 10uM per TA muscle. In a Gastrocnemius muscle, 40 μl in each lobule of the muscle for a more extensive damage (see Fig. 4A).
4 Allow mice to recover fully walking in a 37°C room before you bring them back to the cage. Monitor and provide analgesia accordingly to the animal procedure protocol of the institution for 24 h.

**Fig. 4.**
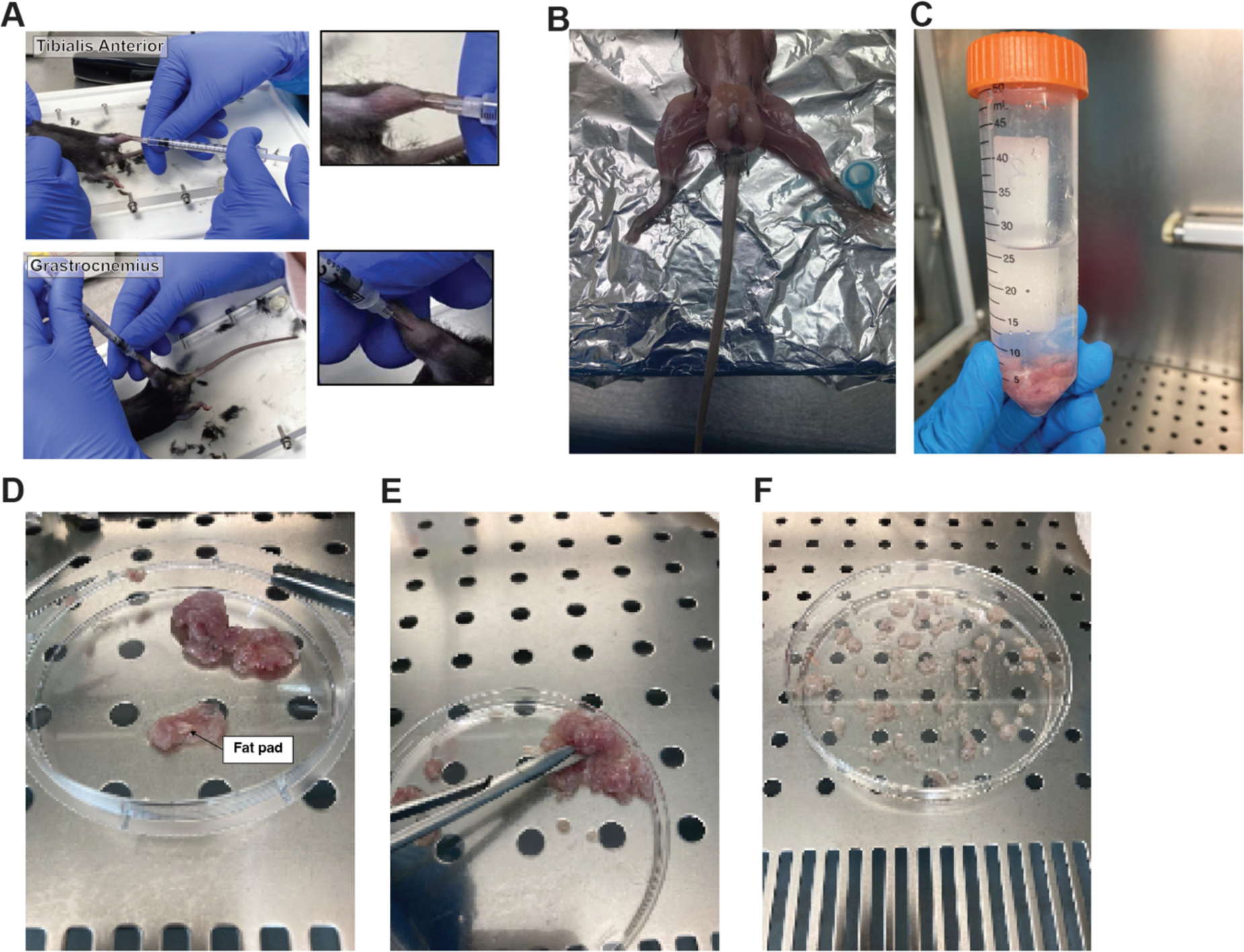
Injured muscle explants preparation. **A** Illustrative pictures of Tibialis Anterior and Gastrocnemius intramuscular injection of cardiotoxin injection. **B** Pre-injured hind limb muscles ready for muscle explants harvest. **C** Hind limb muscles harvested for explants model. **D** Muscle tissue fat and tendons pre-clearing. **E** Hind limb muscles tissue mincing. **F** Representative picture of dropwise muscle dissemination for explant secretion and EVs release.

### Sample collection

5 Four days after injury euthanize mice, accordingly to the animal procedure protocol of the institution.

**Critical step.** Do not proceed to step 6 until there are no signs of reactivity upon gentle pressure on the toes with a surgical forceps.

6 Immobilized the euthanized mice over a Styrofoam lid using pins covered in aluminum foil.
7 Using operating scissors cut the skin perpendicular to the midline around the abdomen and de-glove the skin (see Fig. 4B).
8 Collect all hindlimb muscles from both legs using sterile surgical tools and place them on ice in a 50 ml Falcon tube full of sterile cold PBS. See anatomical images to aid dissection in Fig. 4C.

**Critical step.** Each mouse should be treated as a separate single sample to minimize potential contaminations.

9 Transfer samples into the TC hood.
10 Collect the muscle tissue with a forceps and place all the muscles in a 150 mm petri dish.
11 With the help of surgical scissors and forceps remove all fat depots and tendons to avoid cross-contamination with other sources of EVs.

**Critical step.** Ensure that the large fat pad within the gastrocnemius muscle is removed (Fig. 4D).

12 Once the muscle is clean, cut into small pieces of about 1 mm^3^ (Fig. 4E).
13 Add 50 ml of pre-warmed PBS and transfer the minced muscle with PBS into a 50 ml Falcon tube using a 50 ml pipette.
14 Centrifuge at 300 g for 5 min room temperature in a Benchtop centrifuge using a TX-1000 rotor. Alternatively, another rotor compatible with a 300 g centrifugation step can be used.
15 While tissue is centrifugating, rinse 150 mm Petri dishes, one per mouse, with pre-warmed 37°C FBS EVs-free (to wet the dishes).
16 Aspirate the excess of FBS but do not let the dishes to fully dry.

**Critical step.** Removal of serum is crucial to allow attachment of tissue fragments to culture dish.

17 Once the muscle is pellet down remove PBS carefully with a glass fine pipette connected to the aspirator to avoid muscle loss for aspiration.
18 Rapidly resuspend explants with 10 ml of pre-warmed 37C°C FBS EVs-free.
19 Using a 1000 pipette connected to a tip, spot drops of this mixture on the surface of the wet dishes (Fig. 4F).
20 Lift the side of the dish to remove FBS in excess by with a glass fine pipette connected to the aspirator to avoid muscle loss for aspiration as in step 16.
21 Incubate the dishes on a cell incubator at 37°C, 5% CO_2_ for 2 h to allow explants to adhere to the dish.
22 After 2 h, gently add 30 ml pre-warmed 37°C DMEM medium containing antibiotics with a 25 ml pipette, covering all the muscle tissue.
23 Incubate the dishes on a cell incubator at 37°C, 5% CO_2_ for 2 days.
24 After 2 days, collect medium with a 25 ml pipette into 50 ml Falcon tubes. Discard the muscle tissue into the corresponding biological disposable container.

**Pause point.** At this point, medium can be stored at −80°C prior to proceeding with EV isolation. However, is optimal to continue with the isolation protocol to avoid any loss of samples.

### EVs pre-clarification

25 Centrifuge conditioned medium, obtained in point 24, in a 50 ml Falcon tube at 300 g for 10 min at room temperature to remove floating cells from the mixture. Use a benchtop centrifuge with a swing bucket rotor TX-1000 Thermo Scientific. Alternatively, another rotor compatible with a 300 g centrifugation can be used. If media has been previously frozen, thaw in a water bath at 37°C prior to centrifugation.

**Critical step.** After centrifugation, handle tubes gently to avoid pellet detachment and cross contamination with supernatant.

26 Using a 50 ml pipette, transfer supernatant carefully into a new 50 ml Falcon tube.

**Critical step.** Leave 3 ml of supernatant in the bottom of the Falcon tube to avoid potential aspiration of the pellet and consequent cross-contamination with cells fraction.

27 If pellet is not required as a control, avoid steps 28-29 and go directly to point 30.
28 Resuspend pellet with PBS microparticles-free and spin it once again using same conditions than point 25. This step does not need to be performed under sterile conditions.
29 Aspirate PBS and resuspend pellet in the appropriate buffer (p.e. RIPA buffer for immunoblot) and store accordingly. This step does not need to be performed under sterile conditions.
30 Centrifuge conditioned medium obtained in point 26 at 2500 g for 10 min at room temperature to remove apoptotic bodies from the mixture. Use a benchtop centrifuge with a swing bucket rotor TX-1000 Thermo Scientific. Alternatively, another rotor compatible with a 2500 g centrifugation step can be used.
31 Using a 25 ml pipette transfer supernatant into a sterile 50 ml polycarbonate tube.
32 If the pellet containing apoptotic bodies pellet is required, repeat steps 28-29 but using conditions from step 30. Otherwise continue with point 33.

**Critical step.** Leave 3 ml of supernatant in the bottom of the Falcon tube to avoid potential aspiration of the pellet and consequent cross-contamination with apoptotic bodies fraction.

**Critical step.** Ensure tubes are filled accordingly to manufacturés instructions with the minimal and maximal volume recommended to avoid spills and damage to the tubes.

33 Centrifuge supernatant obtained in point 31 at 20.000 g for 20 min at 4°C to remove microparticles. For this step use a High-speed centrifuge. Different combinations of tubes rotors can be used depending on user availability:

A) Rotor SS34 with Beckman Coulter cat. no. 344058 30 ml tubes.
B) Rotor JA25.50 with Beckman Coulter, cat. no. 357002 50 ml tubes. This is the best option in terms of volume capacity as 8 samples can be centrifuged in one step with bigger capacity tubes.

**Critical step.** Centrifugation conditions are calculated based on the JA25.50 rotor. If a different rotor is used, recalculate the pelleting time taking into account the K factor specific for each rotor [44]. Pelleting time depends on the k factor, a parameter that is specific for each rotor. K factor measures rotor pelleting efficiency at maximum speed. The lower the K factor the shorter running time is going to be at the same speed. Smaller rotors have a lower K factor, as the pathlength of the sample to be pellet down is smaller. This is a critical point as longer running times could compromise the integrity of the vesicle.

34 Using a 25 ml pipette transfer supernatant into a 50 ml FlexyBoy bag (Fig. 3D).
35 If the pellet containing microparticles is required for control purposes, please repeat steps 28-29 but using conditions from step 33. Otherwise continue with point 36.

**Pause point.** At this point, medium can be frozen in the Flexboy bag at −80°C prior to conducting the TFF procedure. However, it is recommended to continue with the isolation protocol for optimal EV recovery.

### Tangential Flow Filtration

36 Connect to the TFF machine the set up circuit (Fig. 3D), using the previously prepared cartridge.
37 Connect the sterile PBS bottle to the auxiliary pump washing port (Fig. 3A).
38 Repeat steps 4-6 from **New cartridge rinsing and sanitation procedure.**
39 Start main pump at 37 ml/min. Make sure there are no bubbles, and all pressures are stable, P_f_ ∼4-6 psi and P_r_ ∼1-2 psi. Run for about 5min.
40 Once pressure is stable, start the automatic backpressure valve at 3 psi to get a TMP of 3 psi.

**Critical step.** Adjust the permeate tubing with a manual clamp (Fig. 3E), to get a positive permeate pressure ∼0.2 psi to dimmish loss of EV material through the cartridge pores due to the TMP.

41 Ensure that shear rate is ∼3000 in the software display, based on the parameters previously setup in points 39-40.
42 Filter until approximately one inch of media remains in the Flexboy bag and stop the pump.
43 Start the step of washing isolated EVs sample in continuous mode as we did in point 11 of **New cartridge rinsing and sanitation procedure.** This means that while washing with PBS, sample is permeating through the cartridge and the volume in the sample bag remains steady.
44 Open the auxiliary pump T-valve and started the auxiliary pump to run at 1.5 ml/min.
45 Re-start main pump at 37 ml/min with TMP valve at 3 psi.

**Critical step.** Maintain the volume in the bag low without allowing it to empty. Flow rate may need to be adjusted (0.5 ml/min-3.0 ml/min depending on sample type). Continue to adjust so that volume is steady, meaning same volume of PBS gets permeated than gets into the sample bag from the auxiliary pump. If the sample is getting too diluted, decrease the flow rate.

46 Tare volume on scale and set up an alarm with the desired amount of washing volume to permeate (use Perm HI STOP for pump to stop at certain volume).

**Critical step.** For muscle explant derived EVs, use double the amount of the starting volume sample due to the high volume of undesired free proteins (e.g. if starting sample is 20 ml, use 40 ml of PBS). However, for Wnt7a HEK293T cell-derived EVs, the same volume of PBS as the sample volume is used to get wash out of the Wnt7a free protein. Adjust the PBS volume, depending on the origin and application of the EVs.

47 When washed volume is reached, stop the pumps, and close the T-valve.
48 Concentrate the sample by re-starting the main pump with a flow of 37 ml/min and permeate the entire volume of the Flexboy bag until bubbles appear in the sample loading tube.

**Critical step.** To have a better control of this step, do not restart the back-pressure valve to set up the TMP at 3 when permeating the sample. The TMP generated by the main pump feed is enough to permeate the remaining volume of the sample. Otherwise, it would be too fast, and sample can be lost.

49 Once all the volume in the Flexboy bag is permeated, stop the main pump, and clamp the Flexboy bag port attached to retentate (left side) using the clamp provided with the Flexboy bag. Then, disconnect the retentate tubing from the clamped Flexboy bag.
50 Take a 15 ml tube to drain the dead volume of the tubing set.
51 Place the retentate tubing into the new tube. Set the flow rate of main pump at the maximum flow, 132 ml/min, and make sure the automatic backpressure valve is open and there is no clamp in the permeate tubing.
52 Start main pump and collect all the retentate fraction into the new tube. This fraction is the EVs fraction ready to use. Volume should be around 5-7 ml (Fig. 3E).

**Pause point.** At this point EVs can be frozen in a Falcon tube at −20°C.

53 For cartridge cleaning, please proceed directly to point 59.

### Size Exclusion Chromatography

54 Once the retentate fraction, that contains the purified EVs, is collected proceed to pellet it down by ultracentrifugation. To reduce the pelleting time is better to use a benchtop ultracentrifuge instead of a regular ultracentrifuge. Transfer the contents of the sample using 1ml pipette to an appropriate tube using:

A) Ultracentrifuge L:90 with rotor 70.1Ti and 10.4 mL polycarbonate bottles with cap (Beckman Coulter, cat. number tubes ref. 355603). Run for 70 min at 32.900 rpm 4°C.
B) Bench ultracentrifuge with rotor TLA.110 and 3.2mL polycarbonate tubes (Beckman Coulter, cat. no. 362305). Run for 30 min at 49,000 rpm 4°C.

**Critical step.** If EVs are for *in vivo* or *in vitro* experiments, is very important to maintain the sterility working in a laminar flow hood and with autoclaved material.

**Critical step.** Depending on the volume, use several smaller 1.5 ml eppendorf tubes with the adaptors instead of 3.2 ml open tubes. The pelleting time is the same.

55 Remove supernatant with a 1000 pipette. Resuspend the pellet in the minimal amount of buffer required to be load into the SEC column, and proceed accordingly to manufacturer’s instructions.
56 Collect the different eluted fractions and proceed to quantification to verify the fraction or fractions that contain EVs of interest, based on the number of particles with a NTA equipment accordingly to manufacturer’s instructions (Figure 5A).
57 Fractions of interest can be combined and pellet down as in point 54 or used directly. If they are required for *in vivo* or *in vitro* bioactivity assays is better to filter them using a syringe microfilter of 0,22µm.
58 Analyze and quantify concentration:

A) Perform immunoblot analysis with different positive and negative EVs markers (Figure 5B).
B) Perform Transmission Electron Microscopy (Figure 5C)

**Fig. 5.**
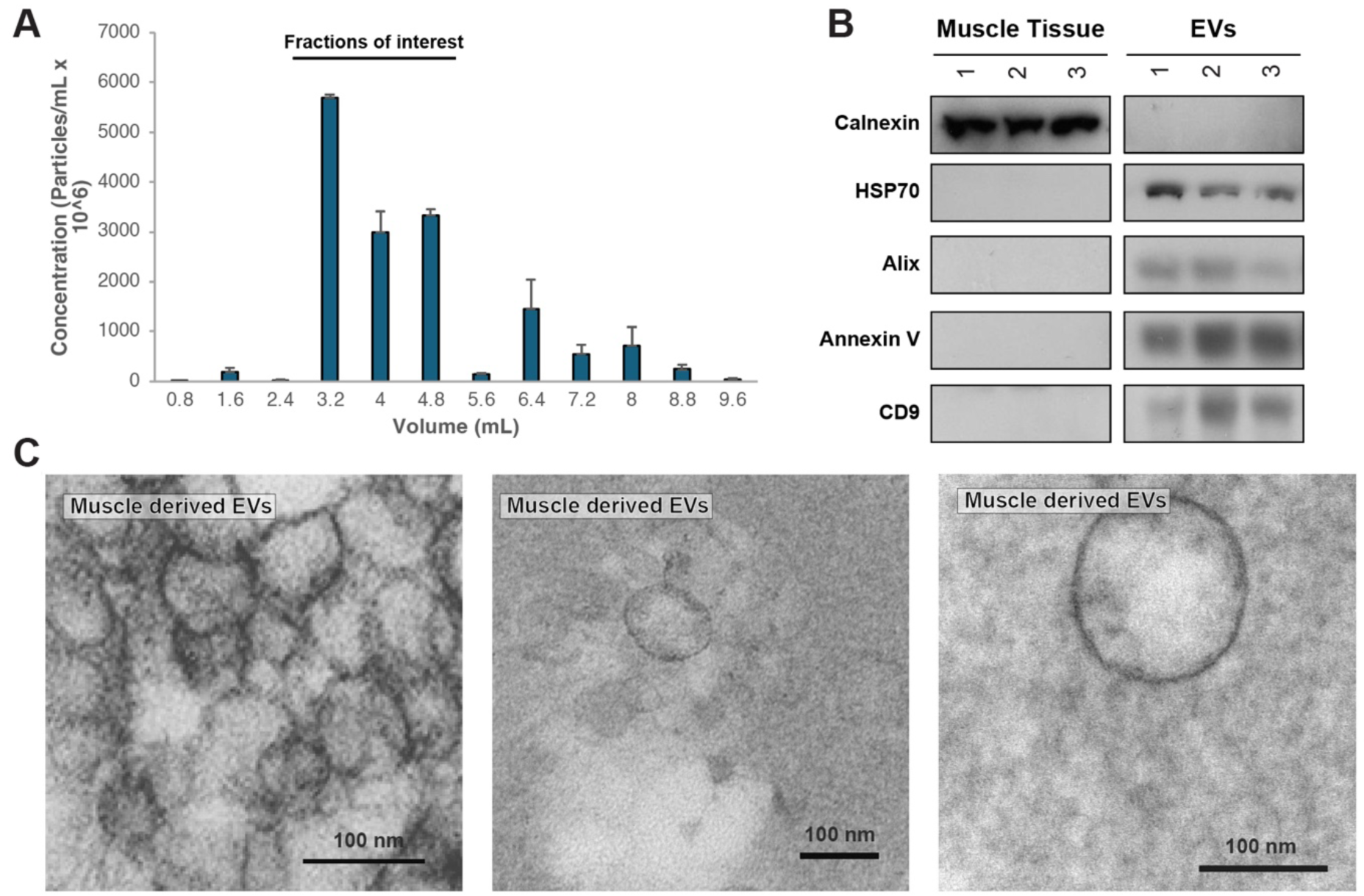
Purified muscle-derived EVs with Tangential Flow Filtration and SEC. **A** Quantification of EVs concentration on the different fractions collected upon SEC. Representative of 3 independent experiments, mean ± s.e.m. **B** Immunoblot analysis showing the expression enrichment of EVs markers on muscle-derived EVs. **C** Representative TEM pictures of isolated EVs derived from new regenerating hindlimb muscles using post-embedding techniques.

### Flush cartridge

59 Change the PBS bottle of the auxiliary pump for the sterile Milli-Q water bottle used in the previous cartridge activation. Screw a plastic cap on the T-valve to cover the open end of the retentate port. Change permeate collection for waste bucket and connect both permeate and retentate into this bucket. Open auxiliary pump and T-valve so circuit is open to Milli-Q water and tubing is not obstructed. Unclamp the retentate port of the Flexboy bag. Make sure that back pressure valve is still opened and main pump is set up at 132 ml/min and there is no clamp in the permeate tubing (Fig. 3E).
60. Start main pump and flush 25 ml of MilliQ water through the system.
61. Stop the main pump. Close the auxiliary pump. Remove the plastic cap on the T-valve and reconnect retentate tubing to Flexboy bag port.
62. Start the auxiliary pump at flow rate of 10 ml/min. Start the main pump at 132 ml/min and flush another 25 ml of MilliQ water with the backpressure valve opened.
63. Stop the pumps.
64. To purify another sample, proceed with step 36. Otherwise continue with point 65.
65. Upon complete of purifications, rinse with NaOH to sanitize the cartridge.
66. Change the auxiliary pump bottle for the 0.5 M NaOH used in the previous cartridge activation.
67. Set up the auxiliary pump for 1-2 ml/min to wash the cartridge in continuous washing mode as in steps 43-46. Rinse with 0.5 M NaOH at main pump flow rate 37 ml/min and with backpressure valve opened. Keep settings until permeate 25-50 ml is complete.

**Critical step.** If media contains red phenol, make sure that the colorant is all washed off and the cartridge is not reddish anymore.

68. Stop the flow and connect retentate to waste. Screw a plastic cap on the T-valve to cover the open end of the retentate port. Connect the sterile PBS bottle to the auxiliary pump. Wash with 50 ml of PBS at maximum speed, 132 ml/min with the back pressure valve open, to flush the remaining NaOH.
69. Stop the flow. Remove the plastic cap on the T-valve and connect retentate flow for recirculation, back to the continued circuit. Initiate the flux with a speed rate of 37 ml/min and TMP 3psi.

**Critical step.** Recirculate 50 ml of PBS and check with pH paper strips that pH is 7 approximately, meaning non traces of NaOH remain inside the circuit that could damage the upcoming sample purification. Otherwise keep recirculating PBS until optimal neutral pH 7 has been reached.

70. Column is ready for use again.

**Critical step.** Keep a record of the number of times the filter has been used and how much volume was filtered. Cartridges can be reused multiple times depending on the type of sample and how concentrated it is. With EVs from HEK293T cells, up to 1L of filtrate can be prepared as long as the cartridge has been washed and stored properly. However, this volume can change depending on the origin of the conditioned media and the conditions of its use.

71. Save and close worksheet, choosing to “End Collection”. Clamp and detach sample bag from circuit, closing tubing ends with plastic caps. Detach other components (e.g. PBS tubing) and close off with plastic caps. It is better to storage the cartridge separately from the tubing and sensors in a plastic bag in a drawer protected from light and high temperatures.

### Troubleshooting

- If pressure is high when instrument is turned on. Turn it off and switch on again.
- If sensors provide a wrong pressure, make the proper tubes have been connected. Color coding the tubes is helpful (e.g. red=permeate, blue=feed and green=retentate).
- Secretion time needs to be optimized and may vary for each type of experiment.

If pressure remains high, the cartridge may be clogged and needs to be washed or changed.

## DISCUSSION

Tissue-derived EVs have recently emerged as ponderable sources of information with a high relevance due to its clinical potential [10,11,38,45]. However, standardized protocols to specifically obtain muscle tissue-derived EVs have not been implemented yet. The standard procedures for tissue-derived EVs are performed with enzymatic digestion procedures to retrieve the EVs within the tissue [38]. Such an approach has the potential to degrade proteins located on the surface of EVs. Finally, EV isolation protocols are typically based on sequential ultracentrifugation protocols [46]. This implies that large volumes cannot be processed, as the rotors have a limited volume specially for ultracentrifugation steps. Conversely, TFF is readily scalable using commercially available cartridges. Moreover, there are pre-setup tables that allow upscaling smaller pilot experiments to adapt to any volume, even for GMP applications [43].

TFF has emerged as a non-ending filtration technique that isolates EVs, retaining the vesicle inside the TFF cartridge within the retentate fraction, and non-EV bound free protein is filtrated through the pores of the cartridge and is collected in the filtrate fraction [39,40]. This is a key point when working with proteins that are secreted in several different modes such as Wnt [29]. Otherwise, EVs products are cross-contaminated with non-EV bound free protein, that could lead to false-positive results.

Overall, using TFF for *in vivo* and *in vitro* down streaming applications is particularly highly recommended. First, TFF is compatible with highly starting volumes as these types of applications usually require large numbers of EVs. This is a very important point when the secreting cells or tissue that do not secrete high numbers of EVs and large volumes are required to achieve desired yields. Second, TFF allows exchange buffer, in this case PBS. Thus, the final product will be already in the actual buffer further required for bioactivity assays. Third, TFF is a gentler procedure, so the characteristic EV double lipid layer keeps intact. Thus, downstream bioactivity applications are not impacted by the loss of EVs morphological integrity.

Importantly, our EV isolation protocol could be used for other types of tissue explants experiments or for the isolation of cell-derived EVs from conditioned media. However, it would be necessary to adjust the secretion and timing conditions accordingly to the experiment. We have used injection of cardiotoxin into the TA muscle to stimulate Wnt7a expression and EV secretion. Other modes of injury or challenges can be contemplated including exercise, disease status, or hormone or drug treatment. Overall, this is a standardized and reproducible protocol to isolate secreted EVs from injured muscle in a timely and efficient manner.

## MATERIAL AND METHODS

### Cell culture

HEK293T cells were obtained from ATCC (CRL-3216) and verified to be free from mycoplasma contamination using the MycoSensor PCR Assay Kit (Agilent Technologies). Cells were cultured as in DMEM (Lonza) supplemented with 10% FBS, 100 U/mL penicillin, 100 U/mL streptomycin and maintained at 37°C in a humidified incubator equilibrated with 5% CO2.

### Conditioned media production for cell derived EVs

Equal numbers of HEK293T cells were seeded Wnt7a plasmid was transfected with linear polyethylenimine (Polysciences), accordingly to manufacturer’s instructions. Later, transfected cells were cultured with 10% FBS EV-depleted [46] in DMEM (Gibco) and maintained at 37°C in a humidified incubator equilibrated with 5% CO2. After 48 h of secretion conditioned media was collected for EVs isolation and samples were purified following the protocol described in this manuscript.

### Pre-embedding immunogold labeling for EVs

TEM on EVs was performed following our previous published protocol [33]. Briefly, EVs specimens were fixed in Karnovskýs fixative for 2 weeks. Fixed EVs pellets were treated with 0.1% sodium borohydride in PBS. EVs were permeabilized with 0.1% Triton X-100 for 10 minutes. After, EVs were blocked in blocking buffer (10% donkey serum + 0.6% gelatin from cold water fish skin in PBS) for 2 h. EVs were incubated with the primary anti-HA antibody (Benthly) for 48 h. Pellets were incubated overnight with the secondary antibody (Jackson ImmunoResearch). Immunogold-labelled EVs were fixed with 2% glutaraldehyde in 0.1 M sodium cacodylate buffer and enhancement was performed with a silver enhancement kit on the immunogold-labelled cells. All samples were post-fixed with 1% osmium tetroxide in 0.1 M sodium cacodylate buffer. Then, samples were dehydrated in increasing concentration of ethanol and infiltrated in Spurr resin overnight at 70°C.. Ultrathin transversal sections (80 nm) were collected onto 200-mesh copper grids and counterstained with 2% aqueous uranyl acetate and with Reynold’s lead citrate. Finally, specimens were observed under a transmission electron microscope (Hitachi 7100, Gatan digital camera). For our analysis, approximately 50 immunoelectron micrographs were examined per sample at at 100,000X and 150,000X magnifications. Importantly, the same protocol was applied to muscle-derived EVs but without immunolabeling in this case. For detailed information about iTEM muscle-derived EVs please refer to [33].

### Immunoblot analysis

Immunoblot analysis was performed as described previously [47] with the following modifications. The lysates from EVs were not clarified by centrifugation. The immunoblot transferring was performed onto PVDF membranes.

### Pre-embedding immunogold labeling for muscle-tissue iTEM

iTEM was performed following our previous published protocol [33]. Briefly, Tibialis Anterior specimens were fixed in Karnovskýs fixative for 2 weeks. After fixation all segments were subsequently washed with 0.1M sodium cacodylate, and treated with 0.1% sodium borohydride in PBS. Samples were permeabilized with 0.1% triton X-100 and blocked with 10% donkey serum + 0.6% fish gelatin. TA samples were incubated with Wnt7a antibody (R&D). After 48 h incubation, segments were rinsed thoroughly with PBS and incubated overnight with the secondary antibody Ultra small (0.8 nm) Gold conjugated (EMS) in blocking buffer at RT. Later, samples were rinsed with 0.1M sodium cacodylate and post-fixed with 2% glutaraldehyde in 0.1 M sodium cacodylate. Pre-embedding enhancement was realized with silver enhancement kit (AURION R-Gent SE-EM, EMS) according to the manufacturer’s instructions. After enhancement, samples were secondly post-fixed with 1% osmium tetroxide in 0.1 M sodium cacodylate buffer. Then, samples were dehydrated in increasing concentration of ethanol and infiltrated in Spurr resin. Ultrathin transversal sections (80 nm) were collected onto 200-mesh copper grids and counterstained with 2% aqueous uranyl acetate and with Reynold’s lead citrate. Finally, specimens were observed under a transmission electron microscope (Hitachi 7100, Gatan digital camera). For our analysis, approximately 50 immunoelectron micrographs were examined per muscle at different magnifications.

## AUTHOR CONTRIBUTIONS

U.G.R: Conceptualization, Methodology, Validation, Formal Analysis, Investigation, Writing-Original draft and Funding acquisition. M.A.R.: Writing-Review&Editing, Supervision, Project administration and Funding acquisition.

## ACKNOWLEDGEMENTS

The authors thank Jennifer Richie for the mice colony management, Yves De Repentigny (OHRI, Canada) for technical help with immunolabeling Transmission Electron Microscopy, Maria Teresa Rojas (Centro Severo Ochoa, Spain) for technical help with Transmission Electron Microscopy, Juan Anguita (CIC bioGUNE) for the legal advice with animal experimentation. The studies from the laboratory of M.A.R. were carried out with support of grants from Defeat Duchenne Canada, the US National Institutes for Health [R01AR044031], the Canadian Institutes for Health Research [FDN-148387; PJT-183804], Ontario Institute for Regenerative Medicine, and the Stem Cell Network.

## REFERENCES

[1] Kalluri R, LeBleu VS. The biology, function, and biomedical applications of exosomes. Science (1979) 2020;367. 10.1126/science.aau6977.

[2] Van Niel G, D’Angelo G, Raposo G. Shedding light on the cell biology of extracellular vesicles. Nat Rev Mol Cell Biol 2018;19:213–28. 10.1038/nrm.2017.125.

[3] Abbott A. FedEx for your cells: this biological delivery service could treat disease. Nature 2023;621:462–4. 10.1038/d41586-023-02906-w.

[4] Herrmann IK, Wood MJA, Fuhrmann G. Extracellular vesicles as a next-generation drug delivery platform. Nat Nanotechnol 2021;16:748–59. 10.1038/s41565-021-00931-2.

[5] Zhao Z, Wijerathne H, Godwin AK, Soper SA. Isolation and analysis methods of extracellular vesicles (EVs). Extracell Vesicles Circ Nucl Acids 2021. 10.20517/evcna.2021.07.

[6] Zhang Q, Jeppesen DK, Higginbotham JN, Franklin JL, Coffey RJ. Comprehensive isolation of extracellular vesicles and nanoparticles. Nat Protoc 2023. 10.1038/s41596-023-00811-0.

[7] Zhang H, Freitas D, Kim HS, Fabijanic K, Li Z, Chen H, et al. Identification of distinct nanoparticles and subsets of extracellular vesicles by asymmetric flow field-flow fractionation. Nat Cell Biol 2018;20:332–43. 10.1038/s41556-018-0040-4.

[8] Théry C, Witwer KW, Aikawa E, Alcaraz MJ, Anderson JD, Andriantsitohaina R, et al. Minimal information for studies of extracellular vesicles 2018 (MISEV2018): a position statement of the International Society for Extracellular Vesicles and update of the MISEV2014 guidelines. J Extracell Vesicles 2018;7. 10.1080/20013078.2018.1535750.

[9] Van Deun J, Mestdagh P, Agostinis P, Akay Ö, Anand S, Anckaert J, et al. EV-TRACK: Transparent reporting and centralizing knowledge in extracellular vesicle research. Nat Methods 2017;14:228–32. 10.1038/nmeth.4185.

[10] Li S, Man Q, Gao X, Lin H, Wang J, Su F, et al. Tissue-derived extracellular vesicles in cancers and non-cancer diseases: Present and future. J Extracell Vesicles 2021;10. 10.1002/jev2.12175.

[11] Qin B, Hu X, Su Z, Zeng X, Ma H, Xiong K. Tissue-derived extracellular vesicles: Research progress from isolation to application. Pathol Res Pract 2021;226:153604. 10.1016/j.prp.2021.153604.

[12] van Niel G, Carter DRF, Clayton A, Lambert DW, Raposo G, Vader P. Challenges and directions in studying cell–cell communication by extracellular vesicles. Nat Rev Mol Cell Biol 2022;23:369–82. 10.1038/s41580-022-00460-3.

[13] Akbar A, Malekian F, Baghban N, Kodam SP, Ullah M. Methodologies to Isolate and Purify Clinical Grade Extracellular Vesicles for Medical Applications. Cells 2022;11:186. 10.3390/cells11020186.

[14] Ji S, Ma P, Cao X, Wang J, Yu X, Luo X, et al. Myoblast-derived exosomes promote the repair and regeneration of injured skeletal muscle in mice. FEBS Open Bio 2022;12:2213–26. 10.1002/2211-5463.13504.

[15] Rome S, Forterre A, Mizgier ML, Bouzakri K. Skeletal Muscle-Released Extracellular Vesicles: State of the Art. Front Physiol 2019;10. 10.3389/fphys.2019.00929.

[16] Yedigaryan L, Sampaolesi M. Extracellular vesicles and Duchenne muscular dystrophy pathology: Modulators of disease progression. Front Physiol 2023;14. 10.3389/fphys.2023.1130063.

[17] Watanabe S, Sudo Y, Makino T, Kimura S, Tomita K, Noguchi M, et al. Skeletal muscle releases extracellular vesicles with distinct protein and microRNA signatures that function in the muscle microenvironment. PNAS Nexus 2022;1. 10.1093/pnasnexus/pgac173.

[18] Wang K, Frey N, Garcia A, Man K, Yang Y, Gualerzi A, et al. Nanotopographical Cues Tune the Therapeutic Potential of Extracellular Vesicles for the Treatment of Aged Skeletal Muscle Injuries. ACS Nano 2023;17:19640–51. 10.1021/acsnano.3c02269.

[19] Ran N, Gao X, Dong X, Li J, Lin C, Geng M, et al. Effects of exosome-mediated delivery of myostatin propeptide on functional recovery of mdx mice. Biomaterials 2020;236. 10.1016/j.biomaterials.2020.119826.

[20] Nakamura Y, Miyaki S, Ishitobi H, Matsuyama S, Nakasa T, Kamei N, et al. Mesenchymal-stem-cell-derived exosomes accelerate skeletal muscle regeneration. FEBS Lett 2015;589:1257–65. 10.1016/j.febslet.2015.03.031.

[21] Aminzadeh MA, Rogers RG, Fournier M, Tobin RE, Guan X, Childers MK, et al. Exosome-Mediated Benefits of Cell Therapy in Mouse and Human Models of Duchenne Muscular Dystrophy. Stem Cell Reports 2018;10. 10.1016/j.stemcr.2018.01.023.

[22] Clevers H, Loh KM, Nusse R. An integral program for tissue renewal and regeneration: Wnt signaling and stem cell control. Science (1979) 2014;346. 10.1126/science.1248012.

[23] von Maltzahn J, Bentzinger CF, Rudnicki MA. Wnt7a-Fzd7 signalling directly activates the Akt/mTOR anabolic growth pathway in skeletal muscle. Nat Cell Biol 2012;14:186–91. 10.1038/ncb2404.

[24] Le Grand F, Jones AE, Seale V, Scimè A, Rudnicki MA. Wnt7a Activates the Planar Cell Polarity Pathway to Drive the Symmetric Expansion of Satellite Stem Cells. Cell Stem Cell 2009;4:535–47. 10.1016/j.stem.2009.03.013.

[25] Bentzinger CF, von Maltzahn J, Dumont NA, Stark DA, Wang YX, Nhan K, et al. Wnt7a stimulates myogenic stem cell motility and engraftment resulting in improved muscle strength. Journal of Cell Biology 2014;205:97–111. 10.1083/jcb.201310035.

[26] von Maltzahn J, Renaud J-MM, Parise G, Rudnicki MA. Wnt7a treatment ameliorates muscular dystrophy. Proc Natl Acad Sci U S A 2012;109:20614–9. 10.1073/pnas.1215765109.

[27] Bentzinger CF, Wang YX, von Maltzahn J, Soleimani VD, Yin H, Rudnicki MA. Fibronectin Regulates Wnt7a Signaling and Satellite Cell Expansion. Cell Stem Cell 2013;12:75–87. 10.1016/j.stem.2012.09.015.

[28] Janda CY, Waghray D, Levin AM, Thomas C, Garcia KC. Structural basis of Wnt recognition by frizzled. Science (1979) 2012;336:59–64. 10.1126/science.1222879.

[29] Mehta S, Hingole S, Chaudhary V. The Emerging Mechanisms of Wnt Secretion and Signaling in Development. Front Cell Dev Biol 2021;9. 10.3389/fcell.2021.714746.

[30] Gross JC, Chaudhary V, Bartscherer K, Boutros M. Active Wnt proteins are secreted on exosomes. Nat Cell Biol 2012;14:1036–45. 10.1038/ncb2574.

[31] Luga V, Zhang L, Viloria-Petit AM, Ogunjimi AA, Inanlou MR, Chiu E, et al. Exosomes mediate stromal mobilization of autocrine Wnt-PCP signaling in breast cancer cell migration. Cell 2012;151:1542–56. 10.1016/j.cell.2012.11.024.

[32] Menck K, Klemm F, Gross JC, Pukrop T, Wenzel D, Binder C. Induction and transport of Wnt 5a during macrophage-induced malignant invasion is mediated by two types of extracellular vesicles. Oncotarget 2013;4:2057–66. 10.18632/oncotarget.1336.

[33] Gurriaran-Rodriguez U, Datzkiw D, Radusky LG, Esper M, Xiao F, Ming H, et al. Wnt binding to Coatomer proteins directs secretion on exosomes independently of palmitoylation. BioRxiv 2023. 10.1101/2023.05.30.542914.

[34] Bartscherer K, Pelte N, Ingelfinger D, Boutros M. Secretion of Wnt Ligands Requires Evi, a Conserved Transmembrane Protein. Cell 2006;125:523–33. 10.1016/j.cell.2006.04.009.

[35] Gurriaran-Rodriguez U, Kodippili K, Datzkiw D, Javandoost E, Xiao F, Teresa Rejas M, et al. Wnt7a is Required for Regeneration of Dystrophic Skeletal Muscle n.d. 10.1101/2024.01.24.577041.

[36] Le Gall L, Ouandaogo ZG, Anakor E, Connolly O, Butler Browne G, Laine J, et al. Optimized method for extraction of exosomes from human primary muscle cells. Skelet Muscle 2020;10:20. 10.1186/s13395-020-00238-1.

[37] Le Bihan MC, Bigot A, Jensen SS, Dennis JL, Rogowska-Wrzesinska A, Lainé J, et al. In-depth analysis of the secretome identifies three major independent secretory pathways in differentiating human myoblasts. J Proteomics 2012;77:344–56. 10.1016/j.jprot.2012.09.008.

[38] Crescitelli R, Lä C, Lö J. Isolation and characterization of extracellular vesicle subpopulations from tissues. Nat Protoc n.d. 10.1038/s41596-020-00466-1.

[39] Visan KS, Lobb RJ, Ham S, Lima LG, Palma C, Edna CPZ, et al. Comparative analysis of tangential flow filtration and ultracentrifugation, both combined with subsequent size exclusion chromatography, for the isolation of small extracellular vesicles. J Extracell Vesicles 2022;11. 10.1002/jev2.12266.

[40] Busatto S, Vilanilam G, Ticer T, Lin W-L, Dickson D, Shapiro S, et al. Tangential Flow Filtration for Highly Efficient Concentration of Extracellular Vesicles from Large Volumes of Fluid. Cells 2018;7:273. 10.3390/cells7120273.

[41] Guo J, Wu C, Lin X, Zhou J, Zhang J, Zheng W, et al. Establishment of a simplified dichotomic size-exclusion chromatography for isolating extracellular vesicles toward clinical applications. J Extracell Vesicles 2021;10. 10.1002/jev2.12145.

[42] Théry C, Amigorena S, Raposo G, Clayton A. Isolation and Characterization of Exosomes from Cell Culture Supernatants and Biological Fluids. Curr Protoc Cell Biol 2006;30. 10.1002/0471143030.cb0322s30.

[43] Musumeci T, Leonardi A, Bonaccorso A, Pignatello R, Puglisi G. Tangential Flow Filtration Technique: An Overview on Nanomedicine Applications 2018:48–60. 10.2174/2211738506666180306160921.

[44] Konoshenko MYu, Lekchnov EA, Vlassov A V., Laktionov PP. Isolation of Extracellular Vesicles: General Methodologies and Latest Trends. Biomed Res Int 2018;2018:1–27. 10.1155/2018/8545347.

[45] Zhi Z, Sun Q, Tang W. Research advances and challenges in tissue-derived extracellular vesicles. Front Mol Biosci 2022;9. 10.3389/fmolb.2022.1036746.

[46] Théry C, Amigorena S, Raposo G, Clayton A. Isolation and Characterization of Exosomes from Cell Culture Supernatants. Curr Protoc Cell Biol 2006;3:3.22.1–3.22.29.

[47] Gurriarán-Rodríguez U, Santos-Zas I, González-Sánchez J, Beiroa D, Moresi V, Mosteiro CS, et al. Action of obestatin in skeletal muscle repair: Stem cell expansion, muscle growth, and microenvironment remodeling. Molecular Therapy 2015;23. 10.1038/mt.2015.40.

